# High germline mutation rates but not extreme population size outbreaks influence genetic diversity in crown-of-thorns sea stars

**DOI:** 10.1101/2023.06.28.546961

**Authors:** Iva Popovic, Lucie A. Bergeron, Yves-Marie Bozec, Ann-Marie Waldvogel, Samantha M. Howitt, Katarina Damjanovic, Frances Patel, Maria G. Cabrera, Gert Wörheide, Sven Uthicke, Cynthia Riginos

## Abstract

Lewontin’s paradox, the observation that levels of genetic diversity (π) among animals do not scale linearly with variation in census population sizes (*N_c_*), is an evolutionary conundrum, where the most extreme mismatches between π and *N_c_* are found for highly abundant marine invertebrates. Yet, whether new mutations influence π relative to extrinsic processes remains unknown for most taxa. Here, we provide the first direct germline mutation rate (*μ*) estimate for a marine invertebrate, using high-coverage (60x) whole-genome sequencing of wild-caught *Acanthaster* cf. *solaris* crown-of-thorns sea stars (Echinodermata). We also provide empirical estimates of adult *N_c_* in Australia’s Great Barrier Reef to jointly examine the determinants of π. Based on direct observations of 63 *de novo* mutations across 14 parent-offspring trios, the *A.* cf. *solaris* mean *μ* was 9.13 x 10^-09^ mutations per-site per-generation (95% CI: 6.51 x 10^-09^ to 1.18 x 10^-08^). This value exceeds estimates for other invertebrates, showing greater concordance with reported vertebrate germline mutation rates. Lower-than-expected *N_e_* (∼70,000-180,000) and low *N_e_*/*N_c_* values (0.0047-0.048) indicated significant genetic drift and weak influences of contemporary population outbreaks on long-term π. Our findings of elevated *μ* and low *N_e_* in *A.* cf. *solaris* may help explain high mutational loads and extreme polymorphism levels observed in some marine invertebrate taxa and are consistent with *μ* evolving in response to *N_e_* (drift-barrier hypothesis). This study advances our understanding of the processes controlling levels of natural genetic variation and provides new data valuable for further testing hypotheses about mutation rate evolution across animal phyla.

## Introduction

Understanding how and why genetic diversity varies among taxa is a longstanding evolutionary puzzle (Lewontin 1974). The neutral theory of evolution predicts that genetic diversity should increase proportionally with effective population size, *N_e_* (Kimura 1971), where the expected level of pairwise genetic diversity (π) at neutral sites reflects the balance between new mutations and loss via genetic drift (π *= 4N_e_μ*, where *μ* is the mutation rate per base pair, per-generation). However, empirical evidence shows that the range of genetic diversity observed in natural populations is far smaller than ranges in census population sizes (*N_c_*) (Lewontin 1974; Leffler et al. 2012; Romiguier et al. 2014). This disparity between π and *N_c_* variance across taxa, known as Lewontin’s paradox, contradicts theoretical expectations and challenges our understanding of the processes controlling levels of natural genetic variation. With increasing evidence that genetic diversity is central to many conservation problems (Hoban et al. 2021; Thompson et al. 2023), including inferences about extinction risks and species responses to environmental change, our ability to understand how diversity levels are maintained and how they relate to census population sizes is critical. Yet, the determinants of genetic diversity and the causative factors underlying Lewontin’s paradox are still debated well into the genomic era (Romiguier et al. 2014; Corbett-Detig et al. 2015; Ellegren and Galtier 2016; Buffalo 2021; Charlesworth and Jensen 2022).

Population genetic studies aiming to solve Lewontin’s paradox suggest that neutral and selective evolutionary processes act in combination to decouple π from census population size (Charlesworth and Jensen 2022) (Figure 1). Historical demographic fluctuations and contemporary dynamics such as repeated extinctions and recolonisations, outbreaks of pest populations or invasive species introductions may result in founder events that substantially reduce *N_e_* (Wright 1938; Kimura 1983). Positive selection (Gillespie 2001) and the indirect effects of background selection and recombination rate variation may also reduce linked neutral diversity to levels lower than predicted by census population size (Charlesworth et al. 1995, Corbett-Detig et al. 2015; Charlesworth and Jensen 2022). Importantly, equilibrium values of π are also a function of the mutation rate scalar, *μ,* which varies by many orders of magnitude across animal taxa (Bergeron et al. 2023). Several hypotheses seek to explain variation in mutation rates across organisms. In particular, the drift-barrier hypothesis predicts that more efficient selection (relative to genetic drift) in large populations leads to maximal fine-tuning of DNA repair and replication mechanisms such that reduced germline mutation rates should evolve in highly abundant taxa (Lynch 2010; Lynch et al. 2016; Wei et al. 2022) (Figure 1). Alternatively, life history traits such as differences in generation time (Bergeron et al. 2023) and reproductive longevity (Thomas et al. 2018; Wang et al. 2022) may be important determinants of mutation rate evolution across animal phyla, whereby longer-lived organisms have higher mutation rates compared to shorter-lived taxa (which also tend to be more abundant). While these predictions may help explain lower than expected π in highly abundant organisms, the extent to which mutation rate variation influences π relative to extrinsic environmental and selective processes remains unknown for most taxa.

**Figure 1.**
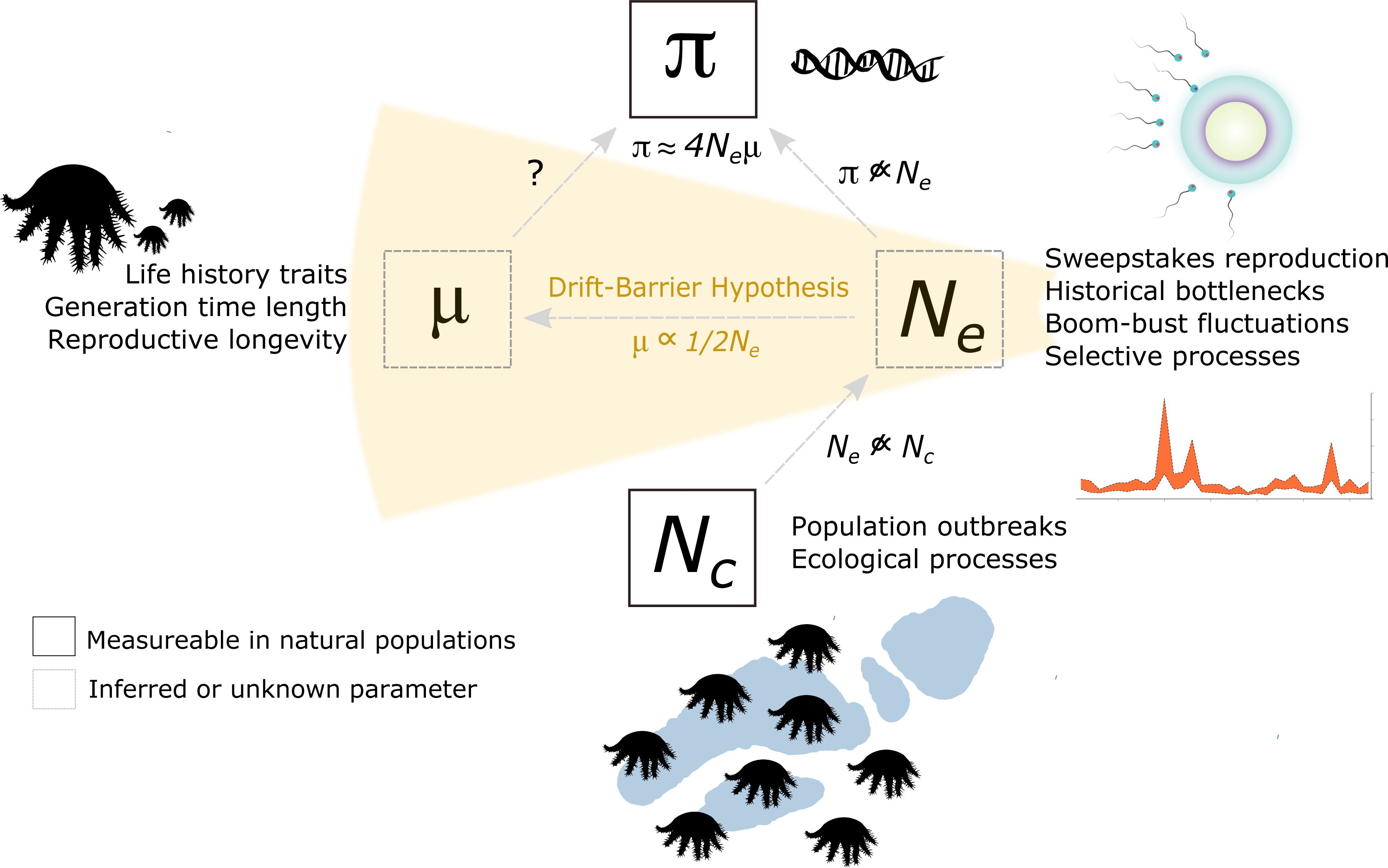
Conceptual diagram illustrating the relationships between evolutionary parameters underlying genetic diversity. Under mutation-drift equilibrium, pairwise genetic diversity (π) reflects the balance between the rate of new mutations (*μ*) and the loss of variation via genetic drift, reflected by effective population size (*N_e_*). In natural populations, π does not increase linearly with *N_e_* and *N_e_* is often much smaller than census population size (*N_c_*)*. μ* is unknown for marine invertebrate taxa, thus the contributions of new mutations to π relative to extrinsic processes remain unknown. A leading explanation for *μ* variation is the drift-barrier hypothesis, which proposes that selection against high *μ* is less efficient in species with small *N_e_.* This leads to an inverse relationship between *μ* and *N_e_* (*μ* ∼1/2*N_e_* for diploid organisms) and the evolution of high *μ* in small *N_e_* populations. Key evolutionary and ecological processes or traits affecting the magnitude of each parameter are shown beside each parameter. Several evolutionary processes act in combination to reduce *N_e_* and thus, constrain π, decoupling it from *N_c_*. This decoupling leads to a disparity between the range of π and *N_c_* variance observed across taxa, known as Lewontin’s paradox. π can be measured in natural populations using DNA sequencing to calculate pairwise differences between sampled individuals and *N_c_* can be approximated most accurately from direct observations of target organisms from field surveys. *μ and N_e_* are inferred parameters from polymorphism data.

Benthically-associated marine organisms have long been recognised for harbouring extreme levels of genetic diversity (Solé-Cava and Thorpe 1991; Leffler et al. 2012; Romiguier et al. 2014). Prolific fecundity and long-lived planktonic larval stages, coupled with few physical barriers to gene flow in the marine environment enable the establishment of large and genetically diverse populations (Gagnaire and Gaggiotti 2016). While the assumption of large population sizes holds true for some marine taxa a possible explanation for high π (e.g., Small et al. 2007), marine invertebrates show some of the most extreme mismatches between π and *N_c_* towards both lower (Hedgecock 1994) and higher relative genetic diversity (Romiguier et al. 2014; Buffalo 2021). High variance in larval recruitment and reproductive success (i.e., sweepstakes reproduction) among broadcast spawning adults may drastically decrease *N_e_* relative to population size (Hedgecock and Pudovkin 2011). Similarly, the ‘boom-bust’ population ecologies of many marine invertebrates, whereby populations rapidly increase in density followed by dramatic declines (Uthicke et al. 2009; Strayer et al. 2017), can decouple *N_e_* from *N_c_* (Figure 1). Specifically, theory suggests that long-term *N_e_* in fluctuating populations is approximated by the harmonic mean and thus bound closer to population sizes during contraction periods (Wright 1938; Motro and Thomson 1982; Nei et al. 1975). From the other perspective, exceptionally high genetic diversity in highly fecund and dispersive species (e.g., tunicates and mussels, Small et al. 2007; Nydam and Harrison 2010; Tsagkogeorge et al. 2012; Romiguier et al. 2014) exceeds expectations based on approximated census population sizes (see Figure 2 in Buffalo et al. 2021). In both cases, decomposing the determinants of π (i.e., *N_e_* and *μ*) and their relationship to *N_c_* is directly relevant for understanding the processes maintaining genetic variation in natural populations (Hare et al. 2011).

**Figure 2.**
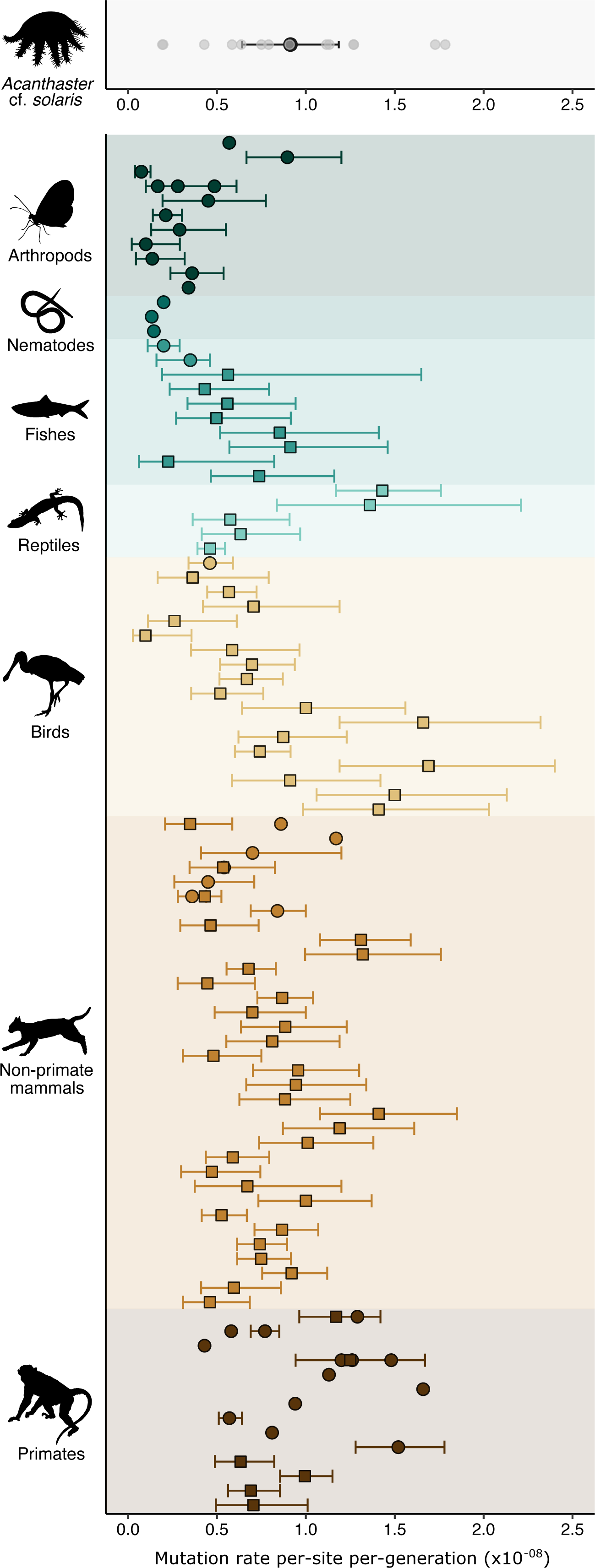
Germline mutation rates inferred in *Acanthaster* cf. *solaris* and other metazoan taxa. Top panel: Per-generation mutation rates for 14 *A.* cf. *solaris* parent-offspring trios. Individual estimates for each trio are shown in grey. The average mutation rate is show in black and 95% confidence intervals are indicated with error bars. Lower panel: Previously published average per-generation mutation rates and 95% confidence intervals for representative metazoan groups. Mutation rates from Bergeron et al. 2023 are represented by squares and estimates from other publications (both pedigree-based and mutation accumulation approaches) are represented by circles. 95% confidence intervals are shown as reported by authors or omitted if not reported. In cases where multiple estimates are available for a single taxon, 95% confidence intervals from the most recent publication are shown. Refer to Supplementary Materials Table S9 for article details. Exceedingly high mutation rate estimates (*μ* > 2.5 x 10^-08^) for *Thamnophis sirtalis* and *Rhea pennata* (Bergeron et al. 2023; Table S9) are omitted for visual purposes only. All silhouettes were sourced from Phylopic (http://phylopic.org).

Delineating whether high contributions of new heritable mutations to genetic diversity could help explain how extreme levels of polymorphism are maintained in some marine invertebrate taxa (e.g., genome-wide silent site π ∼ 8.0% for *Bostrycapulus aculeatus*, Romiguier et al. 2014 and *Ciona savignyi*, Leffler et al. 2012; π ∼ 5.0% for *Ciona intestinalis,* Tsagkogeorga et al. 2012; π ∼ 2.3% for *Crassostrea gigas*, Zhang et al. 2012). Marine invertebrates also show among the highest reported rates of deleterious recessive mutations associated with early mortality (i.e., high mutational load) for animal taxa, including wild populations of mussels, oysters and abalones (e.g., Mallet et al. 1985; Bierne et al. 1998; Harrang et al. 2013; Liu et al. 2006, Plough et al. 2016). Thus, one possible explanation for large mutational loads in marine invertebrates is that their genome-wide mutation rates are substantially higher than terrestrial animals and marine fishes (Plough 2016). Indirect measures of mutational load based on comparisons of synonymous and nonsynonymous diversity from polymorphism data also support elevated genomic mutation rates in the marine invertebrate taxa studied (e.g., Tsagkogeorga et al. 2012; Sauvage et al. 2007; Harrang et al. 2013). Despite the importance of quantifying the contributions of new mutations to genetic diversity, little empirical data exists on germline spontaneous mutation rates for marine organisms. The current knowledge for germline mutation rates in marine metazoans is limited to five fish taxa (Feng et al. 2017; Bergeron et al. 2023), and there are no direct estimates of germline mutation rates for marine invertebrates.

Whole genome sequencing enables us to directly estimate germline mutation rates from a wide range of organisms and is among the most exciting developments in evolutionary biology (e.g., Bergeron et al. 2023). Empirical estimations of mutation rates from parent-offspring pedigrees or mutation accumulation experiments are now available for more than 90 metazoan taxa (Figure 2). Most marine invertebrate life cycles, however, feature a long-lived planktotrophic larval stage that is particularly challenging to raise under controlled conditions. Difficulties in maintaining this critical life stage to metamorphosis and reproduction in captivity have likely precluded mutation rate estimation, as with many non-model and wild populations. Delineating the mechanisms behind Lewontin’s paradox in marine invertebrate phyla is additionally confounded by indirect approaches for estimating census population size. Census population size is typically approximated by individual body size (i.e., as a predictor of population density) and distribution ranges extracted from occurrence data, normally sourced from public databases (e.g., Buffalo 2021; Corebett-Detig et al. 2015). Such estimates are likely to be dramatically underestimated for widespread marine taxa or overestimated for species with either patchy distributions or those comprised of evolutionarily distinct, but morphologically cryptic taxa that are binned together in geographic occurrence data. Knowledge of taxon-specific germline mutation rates would thus enable better understanding of the relationship between π and *N_e_* that is imperative for connecting *N_c_* to population genetic summary statistics as indicators of the eco-evolutionary dynamics of ecologically important marine species (Waples 2022).

In this study, we decompose key evolutionary parameters (*μ*, *N_e_, N_c_*) in a marine invertebrate to investigate the determinants of π and its relationship to population abundance (Figure 1). We present the germline mutation rate, *µ,* for *Acanthaster* cf. *solaris* crown-of-thorns sea stars (CoTS) using high-coverage (60x) whole genome sequencing of pedigree trios of wild-caught parents and early-stage larval offspring, alongside detailed estimates of *N_c_* calculated from 32 years of long-term field monitoring. CoTs are diploid, broadcast spawning asteroid echinoderms that reach sexual maturity at about two years of age (Lucas 1984; Caballes and Pratchett 2014). Although CoTS are native to the Indo-Pacific Ocean, populations can locally reach massive adult densities during cyclical outbreak periods (>1,500 individual ha^-1^; Birkland and Randall 1979; Kayal et al. 2012). As coral predators, CoTS populations cause extensive damage to reef-building corals during outbreak events and are among the leading causes of hard coral cover loss across Australia’s Great Barrier Reef (GBR) (Pratchett 2010; De’Ath et al. 2012; Pratchett et al. 2017). Population outbreaks are followed by rapid declines to population densities several orders of magnitude below outbreaking densities within a few years (density ∼ 3 individual ha^-1^; Uthicke et al. 2022; Rogers et al. 2017). Such boom-bust population fluctuations appear characteristic for echinoderm taxa and are also common to many pests and invasive species. These dynamics are expected to have important consequences on π (Strayer et al. 2017; Chapuis et al. 2009), however empirical datasets against which theoretical expectations can be tested are lacking for natural populations (Charlesworth and Jensen 2022). *A.* cf. *solaris* are one of the most intensely monitored wild marine invertebrate taxa in the world, providing far greater resolution on census population size than possible for most wild marine invertebrates.

Our first aim is to directly quantify *μ* and infer *N_e_*. Given the inverse predicted relationship between *μ* and *N_e_* (i.e., drift-barrier hypothesis), we expect that *μ* is low and similar to values reported for other highly abundant invertebrates, where large, presumed *N_e_* favours the evolution of low mutation rates. Alternatively, if *N_e_* is low, or if elevated genomic mutation rates underly high mutational loads (Plough 2016) and extreme polymorphism levels (e.g., Romiguier et al. 2014) observed in some marine invertebrates, we may expect that *A.* cf. *solaris μ* is high relative to terrestrial animals and marine fish. Similarly, if generation length is an important determinant of mutation rate variation, we expect that the *A.* cf. *solaris μ* exceeds values reported for annual, shorter-lived invertebrate taxa and aligns more closely with mutation rates observed in longer-lived organisms. Our second aim is to clarify the relationship between *N_e_* and *N_c_* to provide insight into the magnitude of genetic drift influencing π in large populations experiencing frequent outbreaks. Our study provides the first germline mutation rate estimate for a marine invertebrate, a crucial basis for interpreting patterns of genetic diversity within species and advances our understanding of the processes controlling levels of natural genetic variation that may help explain extreme cases of Lewontin’s paradox observed in marine invertebrate phyla.

## Methods

### Parental gonad collection and fertilisation

Reproductively mature *A.* cf. *solaris* were collected from the John Brewer Reef (18°38’S, 147°03’E) in the Great Barrier Reef (GBR; November 2020). Collection of gonad tissue, in vitro breeding experiments and larval rearing were performed the National Sea Simulator (SeaSim) Experiment Rooms at the Australian Institute for Marine Sciences (AIMS; Townsville, Queensland) following general procedures outlines in Uthicke et al. 2018. However, in contrast to mixing seven adults of each sex in the spawning, we fertilised individual males and females as unique parental pairs and raised their embryos and larvae for 3-8 days. Refer to Supplementary Materials and Figure S1 for details.

### DNA extraction, genomic library preparation and sequencing

For each parental cross, we selected two larvae in the early-mid bipinnaria stage (day 3-8) or early-mid brachiolaria stage (day 8-9) (Figure S1). DNA from individual larvae was extracted using the QIAGEN Blood and Tissue kit and following modifications for *A.* cf. *solaris* larvae from Doyle et al. (2017). Adult DNA was extracted from gonad tissue using a modified CTAB protocol optimised for marine invertebrate tissue (Panova et al. 2016) and purified with PCR-DX Clean beads. Whole genome libraries were prepared for seven biparental families with two larval offspring per family (to generate 14 mother-father-offspring trios including siblings). We used the NEBnext Ultra II FS DNA library preparation kit following manufacturers specifications, with 10-100ng of input DNA and enzymatic fragmentation for 10 minutes to achieve average insert sizes of 300-400 bp. Final amplification consisted of 4-6 cycles as recommended by the manufacturer. Twenty-eight individually dual-indexed genomic libraries were pooled and sequenced on a single lane of the NovaSeq S4 300 cycle to achieve 60X coverage using 2 x 150 bp paired end reads. Sequencing was performed at the Australian Genome Research Facility.

### Genomic data processing and mapping

FASTQC was used to examine read quality and adapter contamination (http://www.bioinformatics.bbsrc.ac.uk/projects/fastqc). Raw reads were filtered using Trimmomatic. Adapter sequences were removed using the Illuminaclip option in ‘palindrome mode’ and reads were trimmed in 4bp sliding windows with a minimum phred score quality of 20 and a minimum read length of 40 bp. Trimmed reads were mapped to the *A.* cf. *solaris* reference genome (GCF_001949145.1_OKI-Apl_1.0; Hall et al. 2017; NB: the genome of *A.* cf. *solaris* was incorrectly referred to as *A. planci*; see Haszprunar & Spies 2014 and Haszprunar et al. 2017 for details about name assignment of *Acanthaster* species) using the Burrow-Wheeler Aligner (BWA; Li 2011) version 0.7.17 and the MEM algorithm removing reads with mapping quality < 10. The resulting alignment SAM files were converted to indexed and sorted BAM files using Samtools v1.10 (Li et al. 2009). We added read group information to individual BAM files and marked and removed PCR duplicates using picard (http://broadinstitute.github.io/picard/).

### Variant calling and site selection

The bioinformatic pipeline used for calling germline mutations from pedigree samples was initially described in Bergeron et al. (2021) and follows best practices principles outlined in Bergeron et al. 2022. All scripts are available on GitHub (https://github.com/lucieabergeron/germline_mutation_rate). Variant calling was performed in GATK v4.0.7.0 and constrained to 693 scaffolds greater than 10,000 bp. Variants were called for each individual and each scaffold separately using HaplotypeCaller in BP-RESOLUTION mode to retain monomorphic sites and associated quality information. The resulting individual gVCF files were collated into a GenomicsDB for each trio using GenomicsDBImport and joint genotyping was performed for each scaffold using GenotypeGVCF. In parallel, the genotypes for all positions (including monomorphic sites) were retrieved from the GenomicsDB and all scaffolds were gathered using GatherVcfs. We analysed relatedness among all individuals to confirm expected parent-offspring relationships using VCFTOOLS (--relatedness2) (Manichaikul et al. 2010). We then selected SNPs and performed site-filtering to remove SNPs based on the following site-specific parameters: QD < 2.0; FS > 20.0; MQ < 40.0; MQRankSum < −2.0, MQRankSum > 4.0, ReadPosRankSum < −3.0, ReadPosRankSum > 3.0, SOR > 3.0. These parameters consider the quality of a call at a given site (QD), mapping quality (MQ), strand bias (FS and SOR), mapping quality bias between reference and alternative allele (MQRankSum), and position bias within reads (ReadPosRankSum). Variant quality score recalibration (VQSR) was not performed as this step would likely remove rare variants that are putative *de novo* mutations (DNMs) (Bergeron et al. 2021; Bergeron et al. 2022).

### Detecting and filtering of de novo mutations

To estimate the candidate DNMs for each trio, we selected only positions identified as Mendelian violations using GATK SelectVariants (-mendelian-violation), where one of the alleles observed in an offspring is not present in either parent. The following filtering criteria were applied to keep only sites in which: (i) The parents were called as homozygous for the reference allele and the offspring was heterozygous (parental alternative allelic depth per site, AD=0); (ii) all individuals within a trio have GQ > 70, DP > 0.5 * average depth of the individual, and DP < 2 * average depth of the individual, where the average depth was calculated based on the VCF files including every position in the genome with a python script *coverage_python.py*; and (iii) the offspring has an allelic balance (AB) between 0.3 and 0.7 meaning that the number of reads supporting the alternative allele is between 30% and 70% of the total reads at this position (only applied to variant sites). Finally, additional quality control was performed by recalling regions with candidate DNMs with bcftools (version 1.2; Li 2011). Any candidates not jointly detected by GATK and bcftools were classified as False Positive (FP) DNMs.

### Manual curation of de novo mutations

All DNMs and mutation-associated regions were manually inspected using Integrative Genomics Viewer (IGV; Robinson et al. 2011) and the original BAM files to rule out mis-mapping errors, validate homozygous parental genotypes and confirm shared DNMs between siblings. We assigned DNMs as spurious if (i) either parent had a one or more reads supporting the alternative allele; (ii) the DNM site showed violations in minimum depth or allelic balance thresholds in offspring or parents; (iii) the DNM was within 10 bp of an indel; (iv) there was sporadic indel variation or zero coverage zones within 100 bp of the DNM indicating unreliable alignments; (v) there were inconsistencies between parental and offspring genotypes at nearby linked SNPs; (vi) one or more reads supporting the alternative allele were present in any offspring or parents in other trios. As an additional measure to rule out false positive mutations, we compared the remaining DMNs to a large whole-genome dataset of 165 unrelated *A.* cf. *solaris* individuals sampled in the GBR to confirm that sites harbouring DNMs were monomorphic in large populations (Popovic I, unpublished data). For this comparison, we retained singletons and allowed for 50% missing data in the population genomic dataset to capture all reliable polymorphisms.

### Germline mutation rate estimation

To estimate the germline mutation rate per-site per-generation, we estimated the portion of the genome for which we had power to detect candidate DNMs, considering all the sites where mutations could be detected (i.e., number of callable sites) and corrections for the false negative rate. This was done by selecting every position in the VCF files (BP_RESOLUTION output) for which both parents were homozygous for the reference allele and all three individuals passed GQ and DP filters (i.e., GQ > 70, DP > 0.5 * average depth of the individual and DP < 2 * average depth of the individual) (as in Bergeron et al. 2021). A false negative rate was estimated as the proportion of true DNMs that could have been filtered out by sites filters and allelic balance filters applicable to polymorphic positions (Bergeron et al. 2021; Besenbacher et al. 2015). This FNR would be calculated as:

FNR = 1 - ((1-FNR_RP)*(1-FNR_MQRS)*(1-FNR_FS)*(1-FNR_AB)),

where the first 3 parameters are the expected proportion of sites filtered out by the ReadPosRankSum (FNR_RP), MQRankSum (FNR_MQRS), and FS (FNR_FS) site filters, according to a known null distribution. The FNR_AB is an estimation of the proportion of sites that would be filtered out by the allelic balance filter, estimated as: FNR_AB = number of true heterozygous sites outside the allelic balance threshold / number of true heterozygous sites in the offspring. Here, true heterozygous sites are defined as having one parent homozygous for the reference allele (HomRef), the other parent homozygous for the alternative allele (HomAlt), and the offspring are heterozygous.

The mutation rate was then estimated for each trio as:

μ = (nb_DNM – nb_FP)/(2*C*(1-FNR)),

where nb_DNM is the number of de novo mutations, nb_FP is the number of false positive mutations, C is the number of callable sites in the genome and FNR is the false negative rate.

### Parental origins and mutation characteristics

*De novo* mutations were phased to their parental origins using a read-back phasing approach of Maretty et al. (2017) (https://github.com/besenbacher/POOHA) to determine the proportion of male-to-female contributions (α) to detected DNMs. DNMs were classified by mutation type and we determined whether mutations resulting in a change from C to any base were CpG sites. We annotated variants (synonymous, nonsynonymous) and predicted their genomic location (exonic, intronic, intergenic, 5’UTR or 3’ UTR, upstream, downstream) with snpEff v5.1 (Cingolani et al. 2012) according to the *A.* cf. *solaris* reference genome annotations (Hall et al. 2017). We assessed whether the number of DNMs in each annotation category was significantly greater than expected by chance. We determined the expected genomic distribution of annotations based on the total number of polymorphic sites for each trio, and quantified the probability of observing DNMs for each annotation category using a hypergeometric distribution function *phyper()* in R (R Development Core Team, 2022).

### Effective population size (N_e_) estimation

We used the new germline mutation rate and mean nucleotide diversity across 14 parental genomes as input into the Watterson estimator *θ = 4Neμ* (Watterson 1975) to estimate long term *N_e._* We used ANGSD v0.934 (Korneliussen et al. 2014) to calculate mean nucleotide diversity. We estimated folded site frequency spectra using *realSFS* and the *saf2theta* option applying a minimum criteria of mapping quality 30, base quality 30, coverage ≥ 10 reads in 100% of individuals (no missing data). We used the thetastat do_stat option to calculate statistics for each site and in overlapping 50kb windows (10kb step size) across the genome. We obtained genome-wide *θ* estimates by dividing the raw estimates of pairwise theta with the number of sites (nSites) provided by ANGSD. We calculated *θ* with population nucleotide diversity (π) and calculated effective population size as *N_e_ =* π */(4μ)*.

We inferred historical changes in *N_e_* (> 10,000 years ago) of phased parental genomes using Multiple Sequential Markovian coalescent (MSMC2) analysis (Schiffels and Durbin 2014; Schiffels and Wang 2020) and adapted scripts from Github (https://github.com/iracooke/atenuis_wgs_pub). We created a mappability mask for the reference genome using SNPable (http://lh3lh3.users.sourceforge.net/snpable.shtml) to mask genomic regions where sequence reads could not be uniquely mapped. The genome-wide mask file was converted to a bed file using the makeMappabilityMask.py script provided within msmc-tools package (https://github.com/stschiff/msmc-tools). Variant calling was performed using bcftools *mpileup* and bcftools *call* removing sites with mapping quality <30, base quality <30 and indel variants. The bamCaller.py script from the msmc-tools package (https://github.com/stschiff/msmc-tools) was used to produce per-scaffold VCF and mask files for each individual to avoid regions where read coverage is excessively low (<0.5x genome-wide average) or high (>2x genome-wide average). We also retained only scaffolds greater than 1M bp (n=124) representing ∼63% of the genome to reduce computational time. Parental genomes were phased using larval offspring and MSMC input files for each trio were generated using the generate_multihetsep.py script from msmc-tools. MSMC analyses were executed for each pair of phased parental genomes (4 haplotypes) using a single randomly chosen offspring for phasing. We applied the default -p parameter as follows (-p 1*2+25*1+1*2+1*3). A distribution of *N_e_* was obtained for each parental pair applying the mean mutation rate as estimated in this study and a generation time of 2 years (Lucas 1984; Caballes and Pratchett 2014).

To infer *N_e_* on more recent timescales (< 200 generations ago) and to generate *N_e_* estimates that are independent of our inferred *μ*, we use the Genetic Optimisation for *N_e_* estimation (GONE) method (Santiago et al. 2020). For this analysis, we used the 14 parental genomes and removed scaffolds less than 1M bp and variants with minimum quality < 30, genotype quality < 30, mean depth below 10 and above 50. We additionally excluded singletons and removed variants with > 10% missing data, resulting in 4,591,501 SNPs. Files were converted to plink MAP and PED formats to generate input files for GONE. We used the default recombination rate of 1 centimorgans per megabase and set the maximum recombination rate between pairs of analysed loci to 0.01 (hc=0.01) as recommended by the authors (Santiago et al. 2020). Each analysis was performed with 50,000 replicates, and default settings for all other parameters. Because the GONE method considers the compounded effects of genetic drift from all previous generations, we calculated the arithmetic mean *N_e_* between the last 10-80 generations to exclude the most recent and distant generations where estimation may not be reliable (Santiago et al. 2020).

### Census population size (N_c_) estimation using long-term monitoring data

We inferred the size of the contemporary population of *A.* cf. *solaris* adults in the GBR Marine Park using three decades of benthic monitoring data (AIMS 2022) combined with high-resolution (10 m) mapping of the reef geomorphology of the GBR (Roelfsema et al. 2021). Refer to Supplementary Materials for details. The available census data were collected using standardised benthic surveys performed between 1991 and 2022 on a selection of individual reefs. For each surveyed reef, individual counts were converted into an estimate of non-cryptic *A.* cf. *solaris* density (i.e., adult density, where adults are approximately > 15 cm diameter) (Figure S2), and a mean value of reef-level density (individuals km^-2^) was calculated for each annual sample of monitored reefs. A 95% confidence interval of annual mean densities was calculated from 500 pseudo-samples generated by bootstrap for each monitored year. Finally, the confidence limits and the mean of the annual mean densities were multiplied by the total surface (3D) area of the preferred *A.* cf. *solaris* habitat of reef-building corals on the GBR (14,199 km^2^, Supplementary Materials).

## Results

### Germline mutation rate in A. cf. solaris

High coverage sequencing of 14 *A.* cf. *solaris* trios resulted in an average coverage of 59x per individual after mapping (minimum 37x and maximum 91x) (Table S1). Analyses of relatedness validated parent-offspring relationships for each trio (relatedness_phi ∼0.25 between parents and offspring) and low parental relatedness (relatedness_phi < 0.015 between parents) (Table S2). Variant calling in GATK resulted in 7,316,821 SNPs post-filtering (mean across trios) and 596 DNM candidates based on Mendelian violations (Table S3; Table S5). Additional filtering for sites with no reads supporting parental alternative alleles (AD=0) resulted in 246 variants (Table S4; Table S5). Following variant calling with an independent approach, 141 of 246 variants passed our selection criteria as candidate DNMs. We further verified 63 out of these 141 mutations (44.7%) as true positive DNMs based on strict manual validation criteria (Table S6; Table S7). Four false positive mutations were present as low frequency variants in the population genomic dataset spanning the GBR (Table S6). The total number of validated DNMs (n=63) ranged between 1 and 9 mutations per trio (Table S5). Approximately 19.0% (12 of 63) of DNMs were shared among siblings where both shared DNMs passed selection criteria.

The average false positive rate was 42% when considering the independent variant calling approach and 31% for manual validation, resulting in a cumulative false positive rate of 73.0% (range 50%-92% among trios; Table S5). False positives arose largely from the incorrect absence of a heterozygous site in the parents where 1 or more reads supported the alternative allele. The high rate of false heterozygous genotypes could be due to read mapping errors, paralog mismapping or incorrect variant calls associated with reference genome quality (Keightley et al. 2014). We estimated the per-site per-generation mutation rate as the number of observed true positive DMNs out of the total number of callable sites while accounting for the FNR, estimated to be 8.4% (Table S5). The number of callable sites ranged from 196,000,000 to 294,000,000 for each trio, representing 71% of the *A. cf. solaris* genome on average (Table S5), which is comparable to other studies (e.g., ∼80%; Smeds et al. 2016; ∼88% Bergeron et al. 2021). The final estimated mean germline mutation rate averaged across 14 trios was 9.13 x 10^-09^ *de novo* mutations per-site per-generation (95% CI: 6.51 x 10^-09^ to 1.18 x 10^-08^).

### Parental origins and mutation characteristics

We established parental origins for 53/63 DNMs, with 27 maternally and 26 paternally inherited phased DNM across all trios (Table S7; Table S8). The ratio of male-female contributions (α) to germline mutations was 0.96, not statistically distinguishable from 1 (Welch two sample t-test; p=0.50; Figure 3A). There was no effect of family grouping on between group variance (ANOVA; p=0.33; Figure S3). Four phased pairs of DNMs shared among siblings showed no parental bias, however the remaining two pairs of shared DNMs could not be phased to parental origins. Out of 63 DNMs, the number of transitions (47) exceeded transversions (16) as is typically observed in eukaryotes, yielding a transition to transversion ratio (ti/tv) of 2.94 (Figure 3B; Table S7). 15.9% of DNMs (10 of 63) were located CpG sites, two of which were classified as missense mutations (Figure 3B). Twenty-four DNMs occurred within introns (34.9%) and four DNMs within intergenic regions (6.3%), and nine variants (14.2%) fell within coding regions (missense and synonymous variants) (Figure S4; Table S7). There was no significant enrichment of annotation categories based on genome-wide expectations (p>0.05) after corrections for multiple tests.

**Figure 3.**
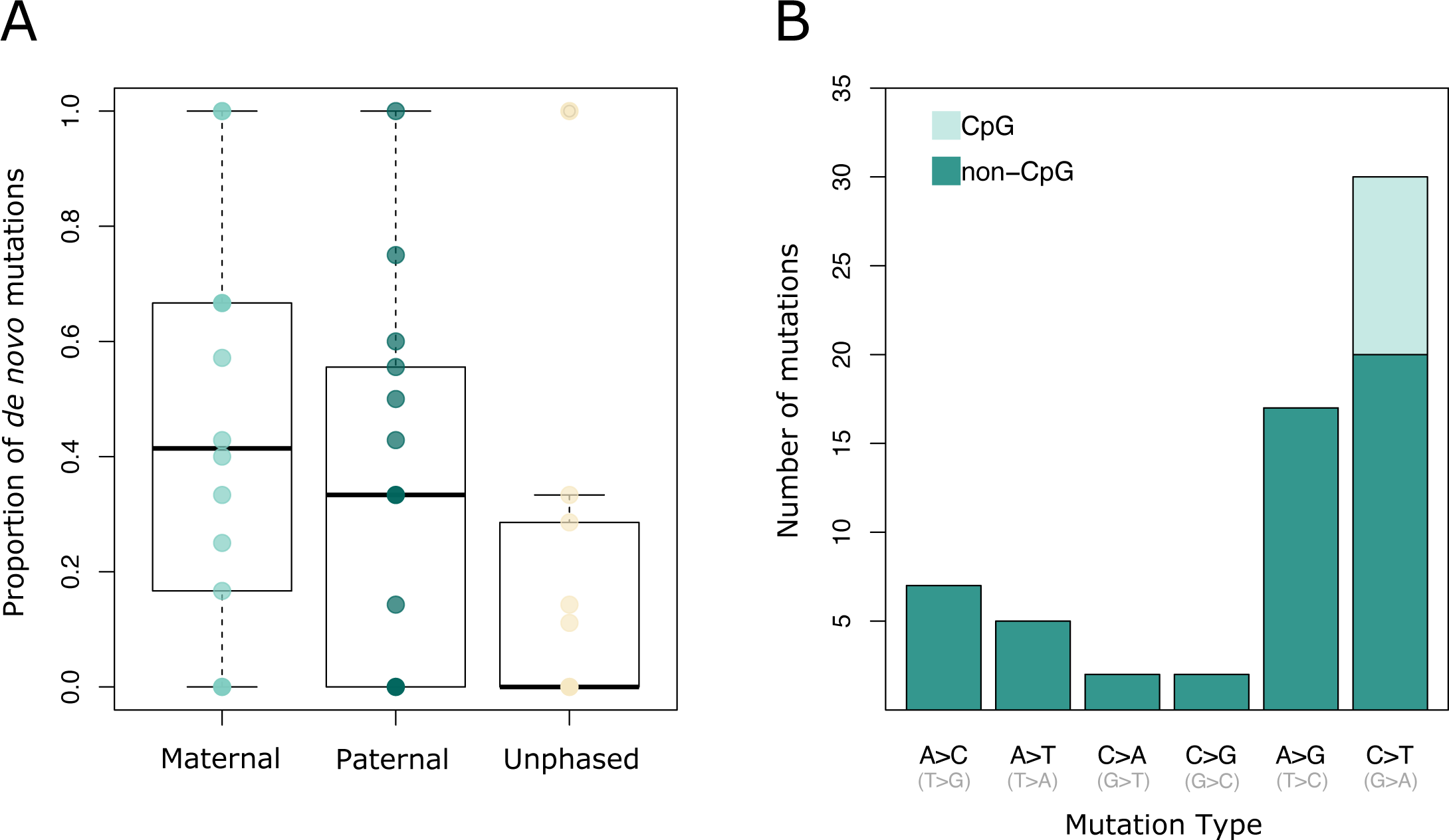
Variation in parental origins and mutational types for *de novo* mutations. (A) Per-trio proportions of *de novo* mutations phased to their parental origins and those with unknown phasing. There was no significant difference in the proportion of maternally and paternally inherited phased *de novo* mutations among trios (Welch two sample t-test; p=0.50); (B) Distribution of *de novo* mutations classified by mutation type, where mutations resulting in a change from C to any base were classified as CpG sites.

### N_e_ and N_c_ estimates

Based on our estimated mutation rate and nucleotide diversity from 14 parental individuals (π = 0.0108), we calculated the long-term *N_e_* (π*/4μ*) of *A.* cf. *solaris* to be 296,000 based on the equilibrium expectations. Historical *N_e_* trajectories inferred by MSMC2 using four parental haplotypes (per larva) showed peak *N_e_* values around 60,000 years ago, followed by a steady decline to a most recent minimum around 20,000 years ago coinciding with the lowest global sea levels during the last glacial maximum (data from Bintanja and Wal 2008; Figure 4). However, more recent *N_e_* estimates < 10,000 years are likely inflated and less reliable because there are only a few coalescent events expected to have occurred on these recent timescales (Schiffels and Durbin 2014). The historical harmonic mean *N_e_* over the past 10,000 to 1 million years was 187,141 and 238,656 over a more recent time period (10,000-100,000 years). Estimates of recent *N_e_* using GONE that are independent of our inferred mutation rate returned a mean *N_e_* of 67,755 between the last 10-80 generations.

**Figure 4.**
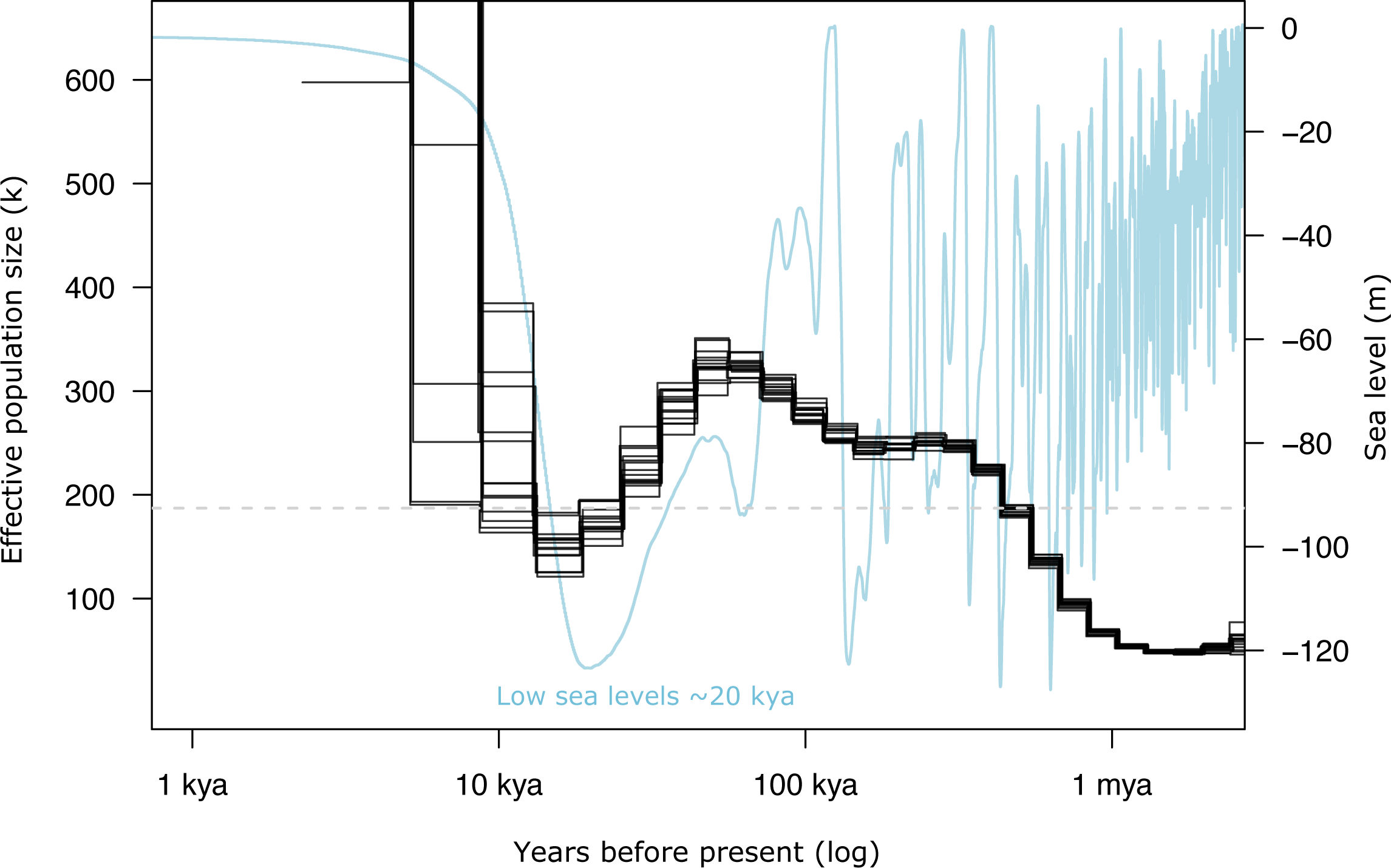
Historical *N_e_* trajectories inferred in MSMC2 applying the *Acanthaster* cf. *solaris* mutation rate and nucleotide diversity from 14 parental genomes (four parental haplotypes per larva). Populations have experienced fluctuations of large magnitudes over the past 10,000 to 1 million years, with peak population sizes around 60,000 years ago and a most recent minimum around 20,000 years ago coinciding with the lowest global sea levels during the late Pleistocene (Bintanja and Wal 2008). The long-term harmonic *N_e_* between the last 10,000 to 1 million years (∼180,000) is indicated by the grey dashed line.

To better understand the ecological factors constraining *N_e_*, we drew upon observational density estimates from 32 years of long-term monitoring surveys to calculate contemporary adult population sizes in the GBR with much higher certainty than is typically feasible for marine invertebrate taxa (e.g., Buffalo 2021). Increased certainty is not only due to the monitoring technique, which allows for the rapid survey of adult populations over large reef areas, but also due to high-resolution mapping enabling reliable estimates of the extent of coral habitat sustaining adult populations of *A.* cf. *solaris*. Between 37 and 136 individual reefs were monitored annually between 1991–2022 across the GBR (Figure 5A). Contemporary *N_c_* estimates across 14,199 km^2^ of coral habitat varied between 6.7 and 14.3 million of non-cryptic individuals (harmonic means of the annual 2.5^th^ and annual 97.5^th^ percentiles of bootstrap distributions, Figure 5B). Population size was 3 to 6 times higher (20–90 million individuals) during outbreaking years (Figure 5B). Considering these *N_c_* confidence intervals, *N_e_/N_c_* ratios ranged between 0.0047-0.048 applying equilibrium *N_e_* (*N_e_/N_c_* = 0.022-0.048), historical *N_e_* (*N_e_/N_c_* = 0.014-0.030) and recent *N_e_* (*N_e_/N_c_* = 0.0047-0.010).

**Figure 5.**
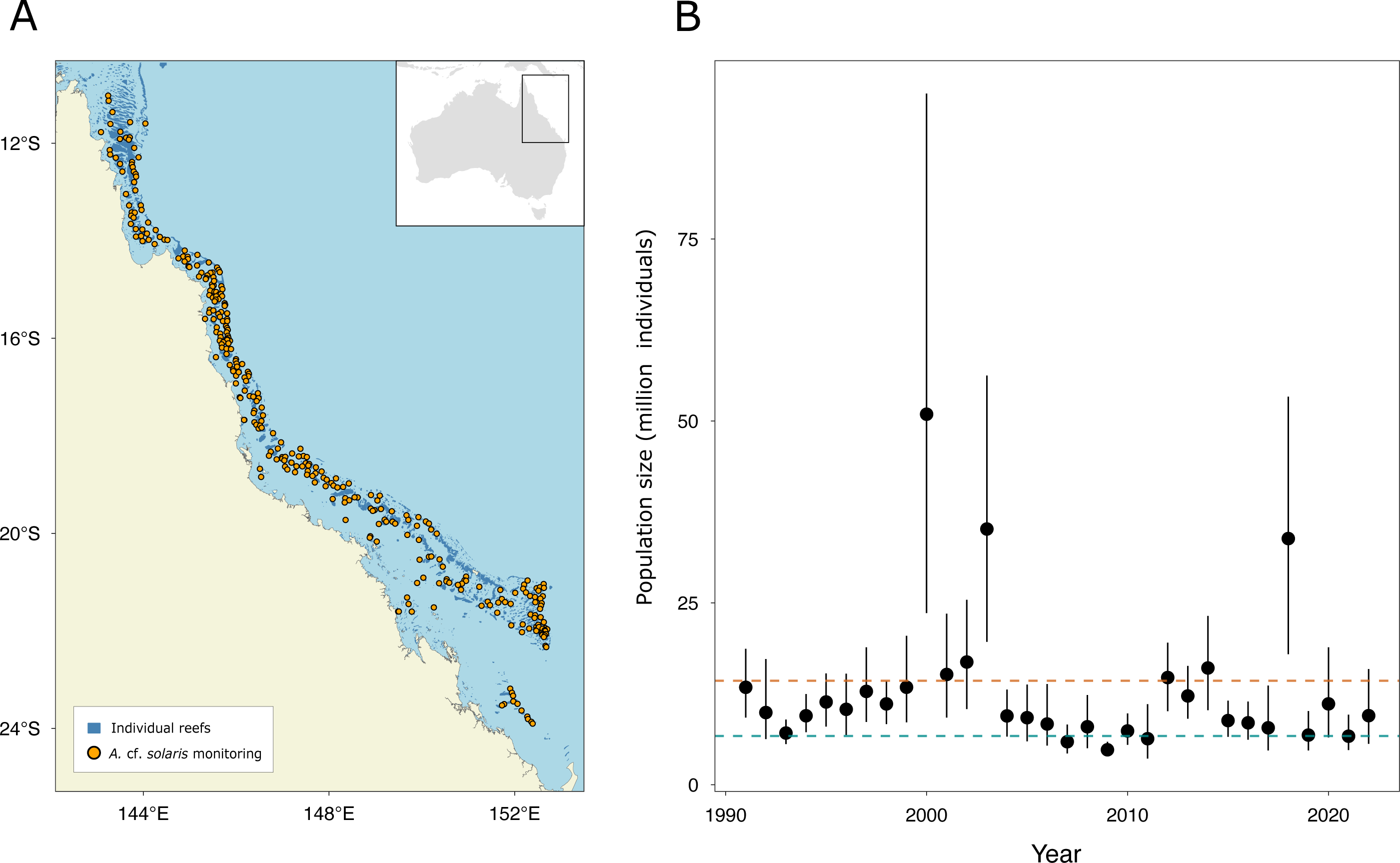
Recent adult *Acanthaster* cf. *solaris* abundance estimates in the Great Barrier Reef (GBR) based on long-term benthic monitoring data. (A) Geographic locations of individual reefs monitored between 1991 and 2022; (B) Contemporary abundance estimates of adult *A.* cf. *solaris* over the entire GBR (3,054 reefs). For each surveyed reef, individual counts were converted into an estimate of *A.* cf. *solaris* density (individuals km^-2^), where mean reef-level density was calculated for each annual sample and a 95% confidence interval of annual mean densities was calculated from 500 pseudo-samples generated by bootstrap for each monitored year. The confidence limits and the mean of the annual mean densities were multiplied by the total surface area of the preferred *A.* cf. *solaris* habitat of reef-building corals on the GBR to calculate census population size. Black dots indicate the average abundance, while vertical lines indicate the extent of the 95% confidence intervals of the 500 mean annual values estimated by bootstrap sampling. The blue and red dashed lines represent, respectively, the harmonic means of the 2.5^th^ (6.7 million) and 97.5^th^ percentiles (14.3 million) of mean annual values over 32 years of monitoring.

## Discussion

Lewontin’s paradox arises from comparisons of genetic diversity (π) and census population sizes (*N_c_*) across the tree of life. Yet, partitioning the determinants of π and characterising species-specific *N_c_* is a difficult challenge, especially for large and fluctuating populations (Ellegren and Galtier 2016; Romiguier et al. 2014; Filatov 2018; Charlesworth and Jensen 2022). In this study, we estimate the germline mutation rate (*μ)* in *Acanthaster* cf. *solaris* crown-of-thorns sea stars and provide empirical estimates of recent adult *N_c_* to interrogate the determinants of π in this species. Based on direct observations of 63 *de novo* mutations across 14 parent-offspring trios, the *A.* cf. *solaris* mean germline mutation rate was 9.13 x 10^-09^ *de novo* mutations per-site per-generation (95% CI: 6.51 x 10^-09^ to 1.18 x 10^-08^). Unexpectedly, our estimate exceeds previously reported values for highly abundant terrestrial and freshwater invertebrates (Figure 2). The *A.* cf. *solaris* germline mutation rate more strongly aligns with values reported for vertebrate taxa, with largely overlapping 95% confidence intervals with both mammals and marine fishes (Figure 2). Using our new estimate of *μ* to infer historical *N_e_* (together with recent *N_e_* estimates that are independent of *μ*), we find that *N_e_* (∼70,000-180,000) is 1-2 orders of magnitude lower than typically reported for highly dispersive marine organisms (e.g., ∼10^6^, Small et al. 2007; Waples 2016) and between 1-2 orders of magnitude lower than *N_c_* estimated from the GBR (Figure 5). Our findings of elevated *μ* and lower than expected *N_e_* in *A.* cf. *solaris* are consistent with the drift-barrier hypothesis, suggesting that short periods (∼2-3 generations) of massive population sizes during outbreak events do not allow for lasting effects of selective fine-tuning of DNA replication machineries to select against high mutation rates. Consistently, we show that *A.* cf. *solaris* exhibits low *N_e_/N_c_* values (0.0047 to 0.048), indicating stronger than expected genetic drift and weak influences of contemporary demographic outbreaks on long-term π. By decomposing key evolutionary parameters (*μ*, *N_e_, N_c_*) and drawing upon multiple sources of data, we advance our understanding about the intrinsic and extrinsic processes shaping π and its relationship to *N_c_*. More broadly, our findings suggest that larger contributions of new mutations may help explain high mutational loads observed in some marine invertebrate taxa (e.g., bivalves; Plough 2016) and high polymorphism levels in marine populations despite demographic declines (Romiguier et al. 2014). Our study also provides new data valuable for further testing hypotheses about the evolution of mutation rates across diverse animal phyla, which are discussed below.

### Low long-term N_e_ and high μ shape genetic diversity in crown-of-thorns sea stars

Using our new estimates of mutation rate and nucleotide diversity from 14 parental genomes (π = 0.0108), the *A.* cf*. solaris* long-term *N_e_* was ∼296,000 based on equilibrium expectations (π*/4μ*). Applying our new mutation rate estimate, reconstructions of historical *N_e_* in MSMC2 returned a lower harmonic mean *N_e_* ∼180,000 over the last 10,000-1 million years. The application of multiple phased genomes allowed higher resolution of more recent population sizes (<100,000 years) than previous coalescent analyses based on single genomes (e.g., Hall et al. 2017). Additionally, the application of a direct mutation rate estimate improved time calibrations revealing significant impacts of historic climactic changes on *A.* cf. *solaris* long-term *N_e_*. We show that *A.* cf. *solaris* have experienced large population fluctuations over the past 1 million years, with the most recent *N_e_* minimum coinciding with the lowest global sea levels during the late Pleistocene (*N_e_* ∼ 150,000; Bintanja and Wal 2008) (Figure 4). Applying the GONE method, which does not rely on *μ* estimates, recent *N_e_* values were even smaller (∼70,000) considering the last 10-80 generations. While *N_e_* estimates in *A.* cf. *solaris* (∼70,000-180,000) are similar or larger than for most terrestrial chordates (e.g., < 10^5^, Bergeron et al. 2023), the *A.* cf. *solaris N_e_* range reported here is 1-2 orders of magnitude lower than for many highly dispersive marine organisms (∼10^6^, Small et al. 2007; Waples 2016; Bergeron et al. 2023), and terrestrial insects (e.g., *Chironomus* non-biting midge ∼2.16-3.95 x 10^6^, Oppold and Pfenninger 2017; ∼1.4 x 10^6^, *Drosophila*, Keightley et al. 2014; ∼2 x 10^6^, *Heliconius*; Kieghtley et al. 2015). This finding adds to empirical evidence that extremely abundant marine taxa, including those with outbreaking tendencies, can sustain low relative effective population sizes, contradicting the broad notion that large marine populations are immune to the effects of genetic drift (Hedgecock 1994; Harrang et al. 2013; Bierne et al. 2016).

From a long-term evolutionary perspective, our findings of lower-than-expected *N_e_* and elevated germline mutation rates in *A.* cf. *solaris* are consistent with the drift-barrier hypothesis as an explanation for mutation rate variation across taxa. The drift-barrier hypothesis predicts that the efficiency of selection on DNA repair and replication machineries is negatively correlated with the strength of genetic drift. Because the power of genetic drift is inversely proportional to the *N_e_* of a species (∼1/2*N_e_* for diploid organisms), high mutation rates are expected to evolve in species with small *N_e,_* such that π is constrained across diverse taxa (Figure 1; Lynch 2010; Sung et al. 2012; Lynch et al. 2016). Indeed, negative correlations between per-generation mutation rates and *N_e_* are evident across the tree of life (e.g., Lynch 2010; Bergeron et al. 2023) and recent work demonstrates that environmental constraints on *N_e_* in prokaryotes can rapidly evolve elevated mutation rates, pointing to evolutionarily labile mutation rates influencing π (Wei et al. 2022). Thus, under the drift-barrier hypothesis, the low *N_e_* in *A.* cf. *solaris* (i.e., similar to terrestrial vertebrates) allows for a high mutation rate to evolve because the ability of selection to maintain high-fidelity replication and repair mechanisms is compromised (relative to the magnitude of genetic drift as reflected by *N_e_*). This observation also helps to explain Lewontin’s paradox by describing the stability of π across diverse phyla as a result of the inverse relationship between *μ* and *N_e_*, rather than a reflection of relatively constant *N_e_* (Lynch 2010).

The low *N_e_* values observed in this study likely reflect historical bottlenecks or *N_c_* minima during non-outbreaking periods. While periods of massive *N_c_* experienced by *A.* cf. *solaris* may increase opportunities for genetic diversity to be reshuffled through phases of high recombination, rare variants are ultimately lost during repeated population bottlenecks, such that genetic diversity can only be recovered through the generational input of new mutations or migration (Wright 1938; Motro and Thomson 1982). Thus, if population size changes occur at a rate higher than coalescent events (that reflect genetic drift), long-term *N_e_* becomes independent of short-term demographic dynamics and contemporary drift (Nordborg and Krone 2002; Sjödin et al. 2005). As such, frequently fluctuating *A* cf. *solaris* populations, are expected to remain far from mutation-drift equilibrium for longer periods of time (Kimura 1964) and a weak relationship is predicted between π and *N_c_* (Romiguier et al. 2014). Consistent with this hypothesis, our calculations of recent adult *N_c_* from long-term monitoring surveys in the GBR resulted in low harmonic mean *N_e_/N_c_* values (0.0047 to 0.048), indicating stronger than expected genetic drift and weak influences of extreme population outbreaks on long-term π. It is important to note that *N_c_* upper bounds reported here were below the distribution of *N_c_* estimates for other echinoderms (∼10^8^-10^12^; Buffalo 2012). Because we only focused on coral habitats as defined by remote-sensing within the GBR (Roelfsema et al. 2021), considering the full-distribution of *A.* cf. *solaris* throughout the Pacific Ocean would have likely resulted in species-wide abundance estimates that are up to 5 x greater than our reported mean *N_c_*. While our *N_e_/N_c_* estimates are conservative for the species-range, we consider them to be precise for the panmictic *A.* cf. *solaris* population within the GBR (Harrison et al. 2017) for which we present genomic data.

In the context of low *N_e_*, larger contributions of new mutations to genetic diversity may thus, in part, explain how moderate polymorphism levels could be maintained in *A.* cf. *solaris,* and may explain extreme polymorphism levels in marine bentho-pelagic taxa more broadly (e.g., tunicates and mussels; Tsagkogeorge et al. 2012; Romiguier et al. 2014). The combination of low *N_e_* and elevated *μ* in *A.* cf. *solaris* (relative to terrestrial invertebrates; Figure 2) could also help reconcile evidence for higher observed mutational loads in marine invertebrates compared to terrestrial taxa and marine fishes (Plough 2016). Because most new mutations are not selectively advantageous, populations with small *N_e_* are expected to hold higher burdens of weakly deleterious mutations due to less efficient purifying selection (Nei 1987). For example, high mutation rates in marine invertebrates have been speculated from evidence for elevated genetic loads in Pacific oysters (e.g., Plough et al. 2016). Similarly, low-frequency deleterious mutations and extreme *dN/dS* ratios in the flat oyster, *Ostrea edulis,* are posited as outcomes of large mutational inputs coupled with small *N_e_* (Harrang et al. 2013). While we provide evidence for a high mutation rate in *A.* cf. *solaris*, the 95% confidence intervals were not distinctly higher than terrestrial animals or marine fishes for which we have germline mutation rate estimates (Figure 2). This result implies that extrinsic processes sustaining low *N_e_* (e.g., sweepstakes reproduction or demographic fluctuations) may have a more predominant role, than mutation rate alone, in shaping genetic load and π in *A.* cf. *solaris* and other marine invertebrate populations. In particular, large variance in reproductive success may be necessary to explain extremely low *N_e_/N_c_* ratios (i.e., < 1%) in wild marine populations of exploited fish and bivalves (Frankham 1995; Vucetich et al. 1997; Waples 2016; Hedgecock 1994; Harrang et al. 2013). High skews in reproductive success may also be relevant for shaping *N_e_* in *A.* cf. *solaris*, especially during non-outbreaking periods when populations can reach densities as low as ∼ 3 individuals ha^-1^ (Uthicke et al. 2022; Rogers et al. 2017). However, testing this hypothesis, and whether reaching known Allee effect thresholds may impose even larger reproductive variance (Eldon et al. 2016) or lower limits to genetic diversity to escape extinction (Romiguier et al. 2014) is challenging to evaluate in marine populations (Waples 2016).

### Alternative hypotheses for mutation rate evolution in A. cf. solaris

There is growing evidence that differences in generation time (e.g., Bergeron et al. 2023) and reproductive longevity (e.g., Thomas et al. 2018; Wang et al. 2022) are important determinants of mutation rate evolution across animal phyla, whereby species with longer generation times have higher germline mutation rates than smaller and shorter-lived organisms (Ohta 1993). This alternative hypothesis could also explain higher *μ* observed in *A.* cf. *solaris* relative to other invertebrate taxa (Figure 2; with the exception of the cyclical parthenogen *Daphnia magna*, 8.9 x 10^-09^, Ho et al. 2020). Compared to annual invertebrates, *A.* cf. *solaris* are longer-lived species (∼8 years in the wild; Pratchett et al. 2017) where most individuals become sexually mature at 2 years of age when individuals become coralivorous (Lucas 1984; Caballes and Pratchett 2014). In captivity, sexual maturity can be delayed up to 6.5 years if juveniles are confined to herbivory (Deaker et al. 2020), implying that generation times in natural populations may be longer depending on the availability of live corals as a food resource. Thus, the mutation rate observed in *A.* cf. *solaris* (values closer to vertebrate taxa; Figure 2) suggests that marine invertebrate taxa may not be an exception to this predictive factor, despite their phylogenetic distance and unique life history characteristics that differentiate them from most chordates, such as extreme fecundity, large census population sizes and bentho-pelagic life cycles (Bierne et al. 2016).

Assigning *de novo* mutations to their parental origin we showed no significant differences in the proportion of male-to-female contributions (α∼1) to germline mutations in *A. solaris*. This result is consistent with similar male and female mutational contributions observed in ectotherm vertebrates (e.g., reptiles mean α=1.5; fishes, mean α = 0.8; Bergeron et al. 2023), in contrast to stronger male biases typically observed in mammals (Kong et al. 2012; Thomas et al. 2018; Wang et al. 2020; Bergeron et al. 2021; Bergeron et al. 2023) which experience larger numbers of mitotic cell divisions during the father’s reproductive lifetime (Crow 2000). This observed variation among ectotherms and mammals may be explained by differences in the process of gametogenesis, whereby seasonally breeding reptiles and fishes produce gametes during limited time periods proceeding mating or spawning activity (Schulz et al. 2010; Gribbin 2011), such that differences in the number of males and female cell divisions are expected to be reduced and α is closer to 1 (e.g., Atlantic herring: Feng et al. 2017). Similar to fishes, in *A.* cf. *solaris* and other echinoderms, highly fecund females produce > 100 million oocytes per season (Babcock et al. 2016) and undergo gametogenesis every spawning period (e.g., November to January; Lucas 1973; Pearson and Endean 1969; Babcock and Mundy 1992). Although little is known about germline formation in sea stars, the source of germline stem cells (gonia) is likely present throughout the year (Bouland and Jangoux 1990; Wessel et al. 2014), while gametes appear to be reabsorbed after the spawning period (Mercier and Hamel 2009; Caballes et al. 2021). The annual renewal of oogonia after the spawning season implies that females replenish germ cells continually for each gametogenesis cycle (Wessel et al. 2014) and that males and females undergo similar numbers of cell divisions throughout their reproductive life spans. In the present study, we show that α estimates for *A.* cf. *solaris* are consistent with theory and previous empirical data for seasonally reproducing animals. The effect of parental age on the observed mutation rate variation among trios (Figure S3) is not clear, however, because unlike other pedigree studies (e.g., Jonsson et al. 2017; Wang et al. 2020), we do not have precise information about the parental age of wild-caught *A.* cf. *solaris*.

### Deep sequencing supports new inferences for non-model bentho-pelagic marine invertebrates

By sequencing 3-day old larvae, we demonstrate that it should be feasible to detect germline mutations in non-model organisms with bentho-pelagic life cycles if pedigreed larvae can be maintained to the earliest developmental stages preceding planktotrophy (e.g., day 3 larvae) - the key stage impeding successful captive rearing of many bentho-pelagic organisms with long-lived larvae. The next main challenge in estimating genomic mutation rates is identifying extremely rare, new mutations against a background of possible sequencing errors (Yoder and Tiley 2021). In the present study, we used deep sequencing (60x) of many trios (>10) and applied several stringent bioinformatic criteria to detect germline mutations with high certainty. First, we applied best practises workflows with conservative quality thresholds (e.g., genotype quality >70; Bergeron et al. 2022) that enabled us to reliably compare the *A.* cf. *solaris* mutation rate with *μ* values from most metazoan estimates (i.e., Bergeron et al. 2023). Second, we mitigated possible limitations imposed by reference genome quality (e.g., Keightley et al. 2014) by visually inspecting BAM files and applying stringent criteria to validate individual *de novo* mutation (DMN) candidates. Third, our additional approach of screening DMN candidates against all sequenced trios and an independent genomic dataset of 165 wild *A.* cf. *solaris* individuals helped ensure that DMNs do not correspond to known alleles segregating in contemporary populations in the GBR. These measures were necessary because resequencing of larvae was not feasible for independent validation. Additionally, our study only examined single base pair mutations and excluded indel variants, multi-nucleotide mutations, copy number variants and larger structural variants that may also be important components of germline mutations as the raw material for evolution (e.g., Sung et al. 2016; Schrider et al. 2011; Oppold and Pfenninger 2017; Thomas et al. 2018; Liu et al. 2017). Although our strict approaches for classifying DNMs may have resulted in underestimations of the *A.* cf. *solaris* mutation rate, we have high certainty in the identified DMNs, and thus confidence in the resulting patterns such as the co-occurrence of DNMs between siblings and trends in parental bias. Our conservative approach therefore further strengthens our main result that mutational rates in *A.* cf. *solaris* crown-of-thorns sea stars are high relative to other measured invertebrate taxa.

## Author contributions

IP, CR, SU and GW conceived the project idea and CR, SU and GW provided funding. SU, MC, FP and KD developed methodologies for larval rearing, collected wild *A*. cf. *solaris*, undertook fertilisation experiments, and generated parent-offspring trios. IP and SH carried out the molecular laboratory work and GW provided guidance on molecular protocols. LB and IP performed bioinformatic analyses to detect and characterise *de novo* mutations. A-MW provided guidance on *de novo* mutation validation and data analysis and assisted with data interpretation. IP performed statistical tests and effective population size analyses. Y-MB developed methodologies for estimating *A.* cf. *solaris* census data and executed the analyses. IP, SH and Y-MB made figures. IP wrote the manuscript draft with all authors contributing to data interpretation, literature review, pitch and editing.

## Acknowledgements

We would like to thank the QRIScould computing cluster at the University of Queensland and the GenomeDK at Aarhus University for providing computational resources for IP and LB, respectively. This work was funded by the Australian Research Council (ARC DP190101593) to CR, SU and GW.

## Data Availability Statement

Raw sequence data from parent-offspring trios will be deposited to the NCBI Short Read Archive. Raw genotype data and relevant code will be made available on The University of Queensland’s Library eSpace.

