## Supplementary Materials for "High germline mutation rates but not extreme population size outbreaks influence genetic diversity in crown-of-thorns sea stars"

*Parental Gonad Collection and Fertilisation*

Adult *A. cf. solaris* were collected from John Brewer Reef (18°38’S, 147°3’E) in the Great Barrier Reef (GBR; November 2020) and transported to the National Sea Simulator (SeaSim) at the Australian Institute for Marine Sciences (AIMS; Townsville, Queensland). Adult *A. cf. solaris* were kept in temperature controlled (26.5 °C) holding tanks supplied with flow-through unfiltered seawater. The experimental room was temperature controlled and set at 28°C ± 0.5°C and contained larval-rearing tanks supplied with filtered seawater (FSW).

Collection of *A. cf. solaris* gonad tissue followed a strict protocol to prevent cross contamination between individuals. After carefully placing the animal on a bench, a small ~1 cm incision was made at the proximal end of each *A. cf. solaris* arm using a sterile scalpel blade and a small amount of gonad was extracted using clean forceps and put on a Petri dish (Figure S1-A). The sex of the individual was visually validated, whereby sperm has a milky appearance and eggs can be seen as individual spheres when the gonad is pressed against the clear surface. While male sea stars were returned to the holding tank, three to four gonadal lobes were removed from each female sea star using forceps and placed in a beaker with 200 mL FSW (Figure S1-B). Each beaker was covered with aluminium foil to prevent contamination with gonads from other individuals. An additional small fragment of gonad tissue was placed in a labelled 2 mL tube with 100% ethanol for genomic analyses.

The extracted female gonads were taken to the experimental room as soon as possible to keep eggs at 28˚C. Male gonads were collected at a later stage (approximately 20 minutes before egg maturation was completed) by making a 1 cm incision as detailed above. The male gonads were placed into a 6-well plate and covered with a lid until the samples were ready to use. In the aquarium room, ovary lobes were washed with FSW over a 500 µm mesh to remove loose eggs and this procedure was repeated several times until no loose eggs were observed (Figure S1-B). To induce maturation and the release of eggs for each female gonad, the beaker was filled with 200 mL of FSW and a 1 mL vial of $10^{-4}$ M 1-Methyl adenine was added and gently mixed with the water. This mixture was left to rest for about 40-70 minutes. During this time, mature eggs dislodged from the gonad and sank to the bottom (Figure S1-C). Once the eggs matured, they were rinsed through a 500 µm mesh to remove unshed eggs and connective tissues and placed in separate beakers for each individual female with about 500 mL FSW.

A sperm solution was made by gently pressing male gonads to release the sperm. For each male, 2 µL of sperm were pipetted in separate scintillation vials and mixed with ~20 mL FSW. Two millilitres of this sperm solution were added to the egg stock solution and carefully mixed using a plunger to guarantee a homogenous solution. After about 5 to 10 minutes, aliquots were taken from the solution to check fertilisation rates under the microscope (Figure S1-D). The egg concentration in the stock solutions were also calculated at this point. The egg stock solution was gently mixed with a plunger to guarantee a homogeneous solution and aliquots were checked under the microscope to validate fertilisation (Figure S1-D).

*Larval Rearing: Culture at 24 hours, 72 hours and 8 days*

Fertilised egg solutions for each cross were transferred to two 2 L jars for a final concentration of 5 eggs x $mL^{-1}$ or about ~10,000 eggs per glass jar. After 24 hours, the larvae were checked under the microscope to ensure that most larvae had reached the gastrula stage (Figure S1-E). An initial sample containing hundreds of larvae was taken and placed in a 2 mL tube with 100% ethanol. The remaining larvae were placed in a new set of clean jars by filtering the culture through individually labelled ~50 µm mesh filters that were placed over a bowl with FSW to prevent the larvae from being damaged. Once all the larvae were filtered, a squirt bottle with FSW was used to push the larvae from the filter into the clean jar, the jars were then filled up with FSW with continued aeration. This procedure was repeated every second day to keep the larvae in good condition.

A second larvae culture sample was taken 72 hours after fertilisation, prior to the start of feeding the larvae. By this time most larvae had reached the early bipinnaria stage (Figure S1-F). On the third day after fertilisation, the early-mid bipinnaria larvae (Figure S1-F-G) started receiving food in the form of a mixture of two algae (*Dunaliella sp* CS-353 and Tisochrysis lutae CS-177) administered twice daily at a ratio of 3:2 and final concentration of 5000 cells $mL^{-1}$. A third sample was taken at day 8 or 9, when most larvae had reached the early brachiolaria to mid-brachiolaria stage (Figure S1-H-I).

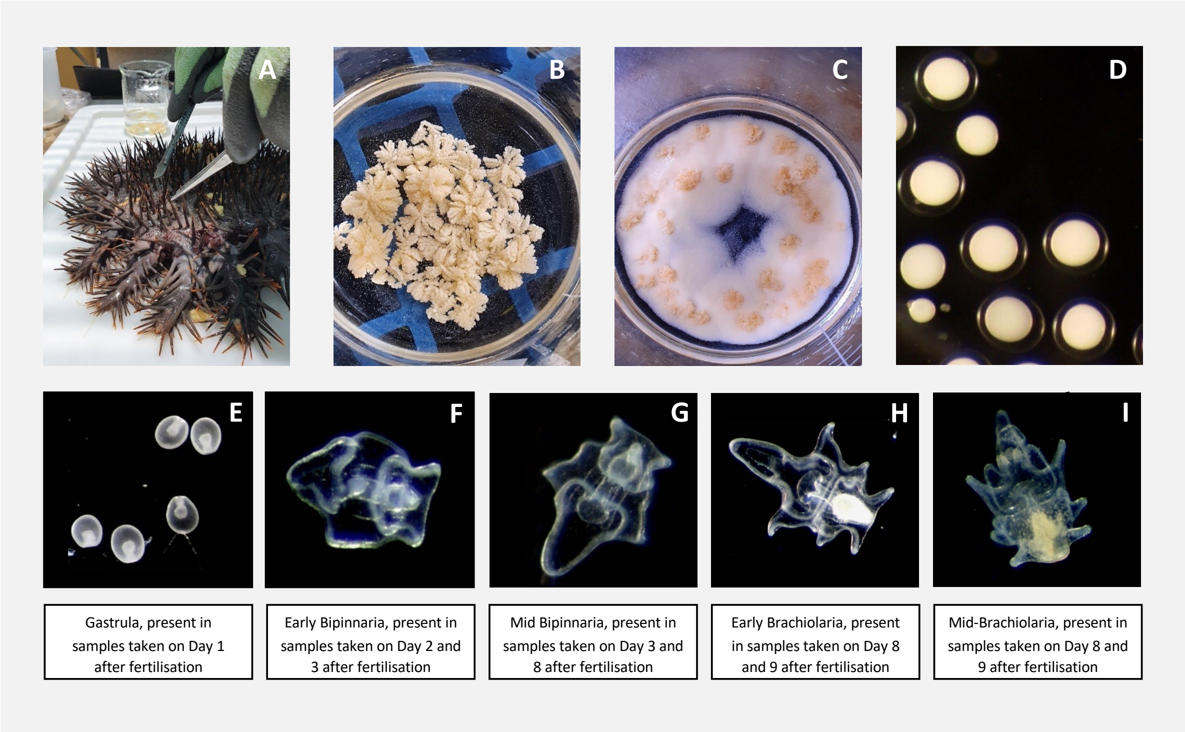


**Figure S1.** Summary of *A. cf. solaris* fertilisation workflow and larval stages. A) *A.* cf. *solaris* adult being dissected to extract gonads; B) Female gonads after rinsing in FSW; C) Mature eggs recently dislodged from the gonad tissue; D) Fertilised eggs under the microscope presenting a clear fertilisation membrane; E-I) Larval developmental stages. For each parental cross, we selected two larvae in the early (F) or mid (G) bipinnaria stage (Day 3-8) and early-mid brachiolaria stage (Day 8-9) (H).

*Crown-of-thorns sea star long-term monitoring data*

Field observations of *A.* cf. *solaris* were obtained from the AIMS Long-Term Monitoring Program (AIMS 2022). The LTMP conducts routine benthic surveys across the Great Barrier Reef Marine Park using the manta tow technique, a standardised procedure allowing for the rapid survey of *A.* cf. *solaris* and corals over large areas of coral reef. The technique consists of towing a snorkel diver at constant speed behind a small boat (Moran et al. 1988, Miller et al. 2009). The observer records from the surface the number of non-cryptic sea stars detected over a period of 2 minutes within a band of approximately 10–12 m along the tow path (depending on boat speed, visibility, reef gradient, distance from the bottom). Individual 2-min tows are maintained parallel to the reef crest and repeated until the entire perimeter of a reef is covered. As such, the manta tow is a cost-effective technique that provides useful information on the broad-scale distribution and temporal dynamics of adult *A.* cf. *solaris* across the GBR (Moran and De'ath 1992a).

Manta tow survey data were available from January 1991 to October 2022. Over this period (32 years), between 37 and 136 individual reefs were surveyed each year across the entire GBR (331 reefs in total). The monitored reefs varied considerably in size, which is reflected by a highly variable number of tows per reef (from 3 to 137, median=42). Tow-level counts were aggregated at the reef level by dividing the total number of *A.* cf. *solaris* recorded for a reef by the number of tows conducted around its perimeter (i.e., mean number of CoTS per tow).

*Calibrated estimates of crown-of-thorns sea star density*

While manta tow surveys are cost-effective, they are also prone to bias due to the low sightability of small individuals and those hidden within the reef matrix (Fernandes et al. 1990). Hence, comparisons with SCUBA swim transects, where the observer carefully inspects the reef matrix in the search of cryptic individuals, have shown that manta tows consistently undercount available sea stars (Fernandes et al. 1990, Moran and De'ath 1992a). Comparing manta tow counts and SCUBA swim counts performed over the same area, Moran and De'ath (1992a) obtained a strong relationship between the two abundance estimates (Figure S2) and argued that, once calibrated, manta tow counts can provide unbiased estimates of SCUBA swim counts. Using their manta tow and SCUBA swim count data, we refine the calibration model and generalise it to predict the density of *A.* cf. *solaris* from the mean number of CoTS per tow recorded on any reef.

Similar to Moran and De'ath (1992a), we performed a linear regression of SCUBA swim counts (SSCs, response) on manta tow counts (MTCs, predictor) after a cube-root-transformation of both variables expressed as per-tow basis (200×12 m, the dimension of a SCUBA transect, roughly equivalent to a 2-min tow search path). Hence, a variable number of tows/transects underlie the reported counts and were used as weights as in the original regression, thus leading to the same model ($R^2$=0.913, Figure. S2, see also equation in Moran and De'ath 1992b):

$$SSC^{1/3} = 0.8071 + 1.2008\ (MTC)^{1/3}$$

The predicted *A.* cf. *solaris* density (D) expressed in starfish.km$^{-2}$ is thus obtained with:

$$D = (10^6 / 2400) \times (0.8071 + 1.2008\,(MTC)^{1/3})^3$$

The uncertainty around density estimates decreases with the number of conducted tows. We can generate a normally distributed noise around D to reflect the variability of reef-level predictions given the number of tows (N) that were required to survey the perimeter of a reef (**Fig. S1**). This is achieved using the R function *rnorm*:

D* = rnorm(mean=D, sd=sqrt(se.fit^2+ residual.scale^2)/sqrt(N))

where D* is a stochastic prediction of reef-level *A.* cf. *solaris* density, random se.fit is the standard error of the mean prediction (D) and residual.scale the standard deviation of the residuals. It is important to note that SCUBA swim searches are not 100% bias-free (underestimating 'true' density by ~11%, Fernandes et al. 1990) so that these predictions cannot be considered as absolute estimates of *A.* cf. *solaris* abundance. However, one can reasonably assume they provide accurate estimates of the density of diurnal (>15 cm), non-cryptic adult sea stars (Moran and De'ath 1992a).

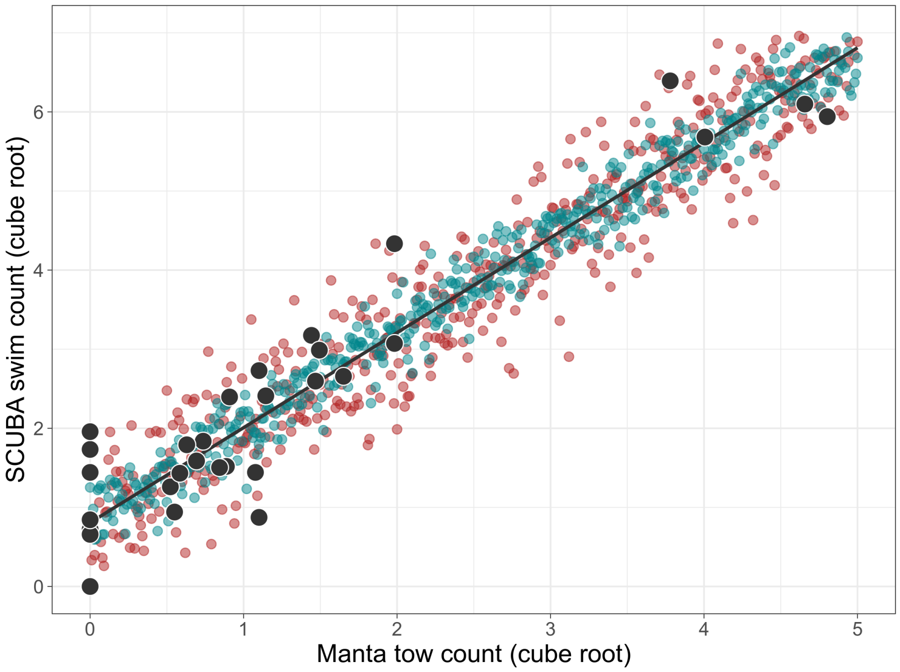


**Figure S2.** Relationship between manta tow counts and SCUBA swim counts from Moran and De'ath (1992a). The observed data (black dots) are expressed as a per-tow basis (200×12 m, with 2 to 15 tows/transects supporting each observation). The regression model is used to generate deterministic predictions of SCUBA swim counts (line) or stochastic predictions function of the number of tows conducted in a given area (e.g., the perimeter of a reef), illustrated here with 5 (red dots) and 25 tows (green dots). Increasing the number of tows decreases the dispersion around the deterministic model.

*Estimates of reef area and A. cf. solaris habitat*

The area of *A.* cf. *solaris* habitat was estimated based on recent high-resolution (10-m) mapping of the GBR geomorphology and substrate type (Roelfsema et al. 2021). This mapping product characterises the geomorphic zonation of reefs to a depth of 20 m using a specific classification of physical attributes derived from remote-sensing data (sub-surface reflectance, bathymetry, slope angle) and wave modelling. Reef geomorphic zonation is classified into 10 nominal categories defined by expert knowledge and validated with in situ observations. The extent of each geomorphic category is given as 3D surface area calculated from the bathymetric profile, thus providing a more accurate estimate of the actual area of reef habitats. The mapping was initially available for 2,164 offshore reefs of the GBR Marine Park (Roelfsema et al. 2021) and was latter extended to include 890 fringing and nearshore reefs (Castro-Sanguino et al. 2023).

As representative *A.* cf. *solaris* habitats, we only considered geomorphic categories that are predominantly covered by consolidated hard substrate, which is more suitable for coral colonisation. According to Roelfsema et al. (2021), there are 4 geomorphic categories that are representative of significant 'coral habitat': 'outer reef flat', 'reef slope', 'reef crest', and 'shelter reef slope'. These habitats are likely to support the greatest share of adult *A.* cf. *solaris* populations, as they provide optimal conditions for abundant shelter and food source. The cumulative 3D area of these 4 geomorphic categories across the 3,054 individual reefs amounts to 14,199 $km^2$ (47.6% of all geomorphic 3D areas). We note that *A.* cf. *solaris* can also be found in habitats deeper than 20 m, and the GBR exhibits significant reef areas below this depth (Harris et al. 2013). However, the extent of suitable habitat for corals appears relatively limited below 20 m (1/3 of the deep-water reef habitat, Beaman 2019), especially for tabular *Acropora* corals which are the preferred prey of *A.* cf. *solaris*. High sea star densities are typically observed in areas of rich coral cover, which are usually found around 10-15 m. Thus, reefal areas deeper than 20 m may be considered as marginal habitats for *A.* cf. *solaris*, unlikely to support outbreaking densities (Beaman 2019).

It is important to note that the present definitions of *A.* cf. *solaris* habitats include inshore and outer reef environments where outbreaking *A.* cf. *solaris* densities are less common compared to mid-shelf reefs (Vanhatalo et al. 2017). While defining *A.* cf. *solaris* habitats accurately is challenging, our two estimates of suitable area for *A.* cf. *solaris* (coral-suitable vs. all geomorphic habitats across 3,054 reefs) can be considered as reasonable bounds (14,199 – 29,827 km$^2$) for the potential extent of significant adult *A.* cf. *solaris* colonisation across the GBR.

*Bootstrap re-sampling and confidence limits*

We define a sample as the collection of all the reef-level MTCs obtained in any given year *y*, which corresponds to the individual reefs $N_y$ sampled by manta tow during that year. Each reef-level MTC is associated to a number of tows ($N_{tows}$) conducted along the reef's perimeter. For each year of monitoring, 500 replicate samples were generated by randomly drawing $N_y$ reef-level MTCs with replacement from the corresponding sample. Within each bootstrap sample, a stochastic prediction of reef-level density D* was generated from each drawn value of reef-level MTC and associated $N_{tows}$ using the calibration model. In doing so, we introduced some variability around density predictions that reflects the uncertainty in detecting *A.* cf. *solaris* from manta tows (consistent with the calibration model). Finally, we averaged the $N_y$ density predictions to estimate the mean reef-level density of non-cryptic *A.* cf. *solaris* per bootstrap sample. This resulted in a distribution of 500 estimates of mean reef-level density for every year of monitoring.

In a second step, mean reef-level densities were multiplied by the 3D area of the total *A.* cf. *solaris* habitat across all 3,054 reefs. The 2.5$^{th}$ and 97.5$^{th}$ percentiles of the resulting distributions were calculated to produce 95% confidence intervals of the annual mean population size of noncryptic *A.* cf. *solaris* between 1991 – 2022.

*Additional Supplementary Figures*

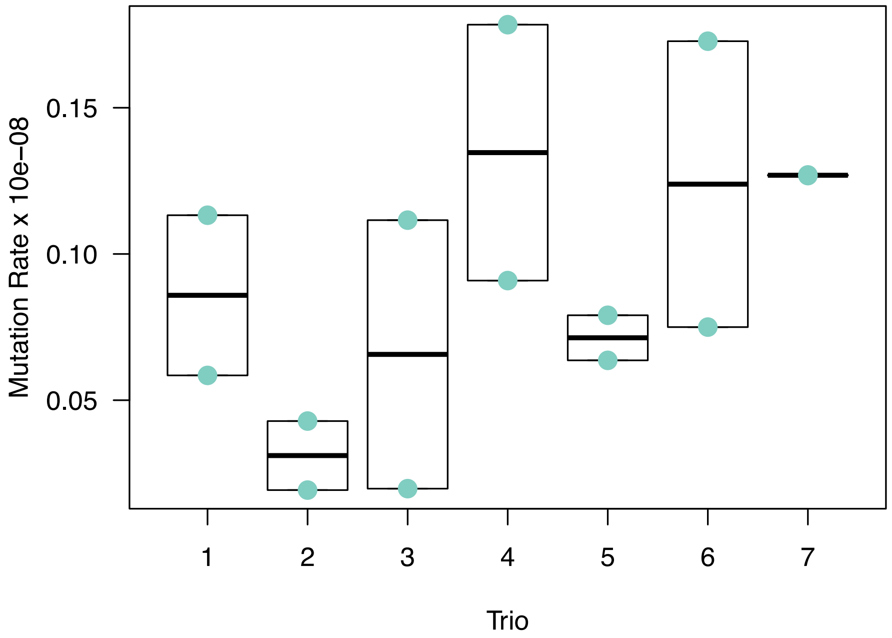


**Figure S3.** Mutation rate estimates for 14 parent-offspring trios grouped by family. There was no effect of family grouping on between group variance (ANOVA; p=0.33). Mutation rate data points for Trio 7 siblings are overlayed.

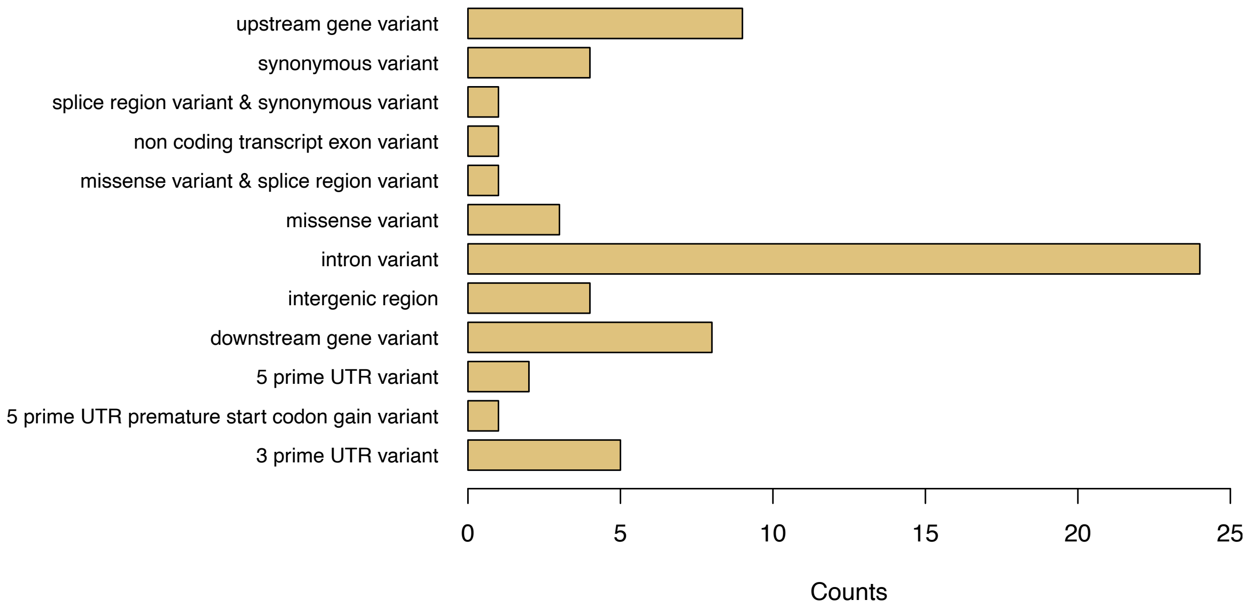


**Figure S4.** Annotated variants (synonymous, nonsynonymous) and predicted their genomic locations according to the *Acanthaster* cf. *solaris* reference genome annotations (Hall et al. 2017). There was no significant enrichment of annotation categories based on genome-wide expectations (p>0.05) after corrections for multiple tests.
